## Supplementary Figures and Description of Supplementary Videos for "SpeedyTrack: Direct microsecond wide-field single-molecule tracking and super-resolution mapping via CCD vertical shift"

### SUPPLEMENTAL INFORMATION

### Supplementary Figures

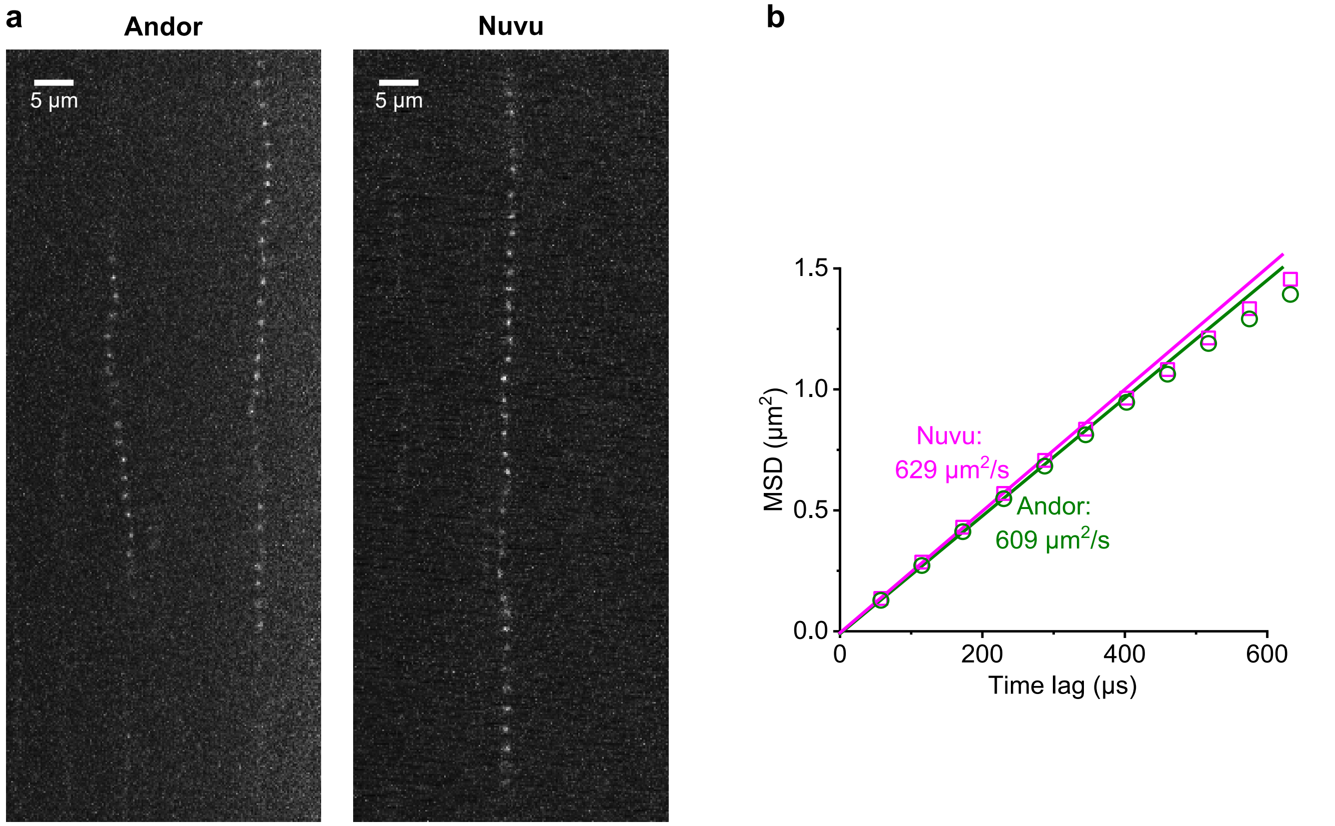


**Supplementary Fig. 1.** Comparison of SpeedyTrack results acquired with two different commercial EM-CCDs for Cy3B diffusing in methanol. (**a**) Example SpeedyTrack raw frames from the Andor (left) and Nuvu (right) EM-CCDs. Both measurements used an exposure time of 50 μs and a vertical shift time of 7.5 μs for 15 rows at each timepoint. (**b**) Data points: MSD vs. time lag calculated from pooled trajectories obtained by the two cameras. Lines: Linear fits to the first 4 data points, yielding diffusion coefficients of 609 and 629 $\mu m^2/s$, respectively.

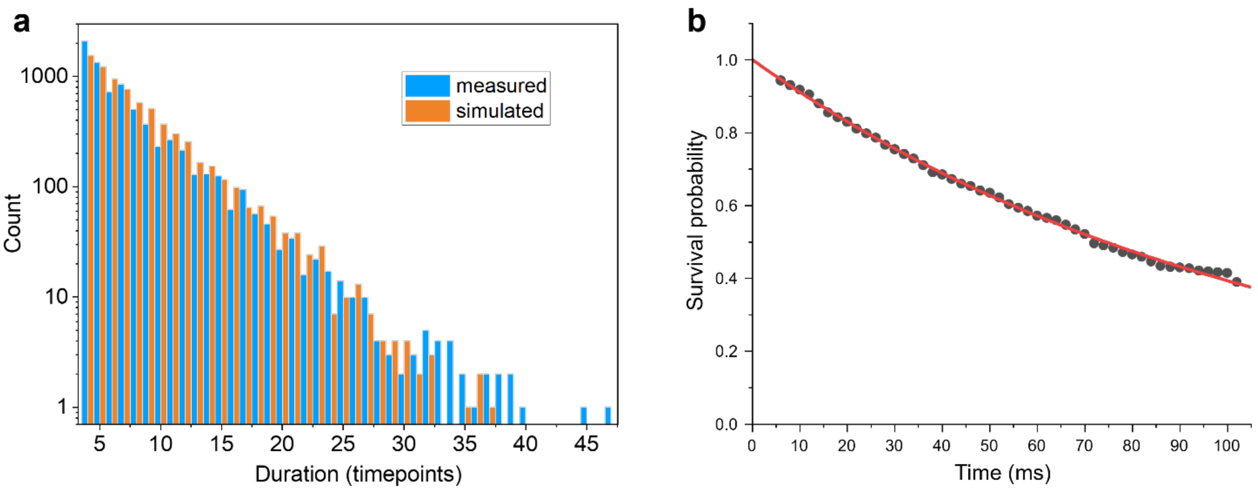


**Supplementary Fig. 2.** Trajectory lengths are limited by diffusion out of the focal plane. (**a**) Distribution of durations for single-molecule trajectories collected by SpeedyTrack for carbonic anhydrase (blue; same as **Fig. 2e**) versus the expected distribution (orange) limited by diffusion out of the focal plane, simulated for the same total number of tracks at $D = 90$ µm$^2$/s and a focal depth of 0.8 µm. Each timepoint is 307.5 µs. (**b**) Photobleaching assay. Avidin (Sigma A9275) was labeled at 0.03 Cy3B per avidin tetramer and then sparsely immobilized on the coverslip to enable single-molecule imaging. To record long single-molecule time traces, SpeedyTrack was performed with an extended exposure time of 2 ms per timepoint under a fixed excitation power of 280 mW (~22 kW/cm$^2$) typical of our single-molecule tracking experiments. Data points: Kaplan-Meier estimator of single-molecule fluorescence survival probability based on the observed track lengths for molecules present at the beginning of the SpeedyTrack frame. Line: fit to an exponential decay, yielding a bleaching time constant of 110 ms (bleaching rate of 9 s$^{-1}$). This value is substantially longer than the typically <10-ms durations of single-molecule tracks in our data.

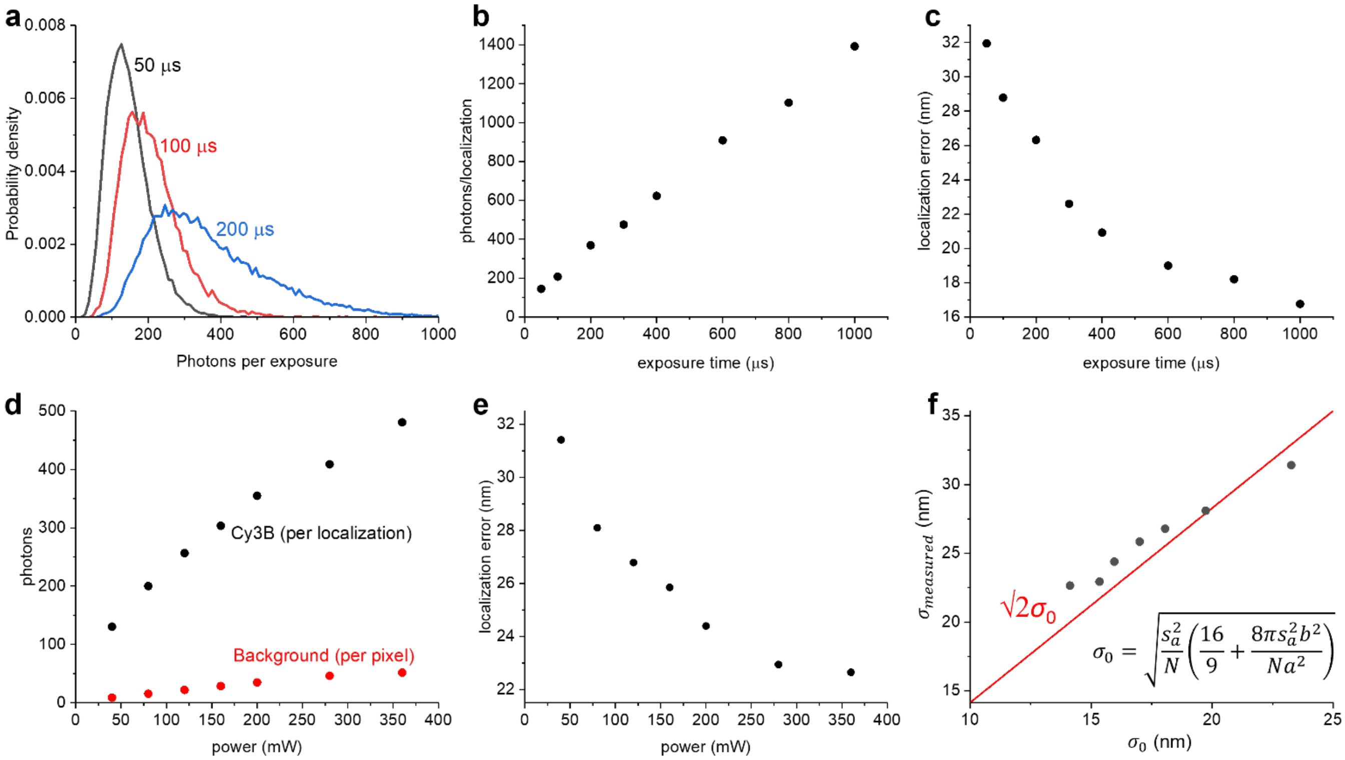


**Supplementary Fig. 3.** Single-molecule photon counts and localization precision of SpeedyTrack on immobilized Cy3B. Avidin (Sigma A9275) was labeled at ~0.4 Cy3B per avidin tetramer and incubated on an acid-washed glass coverslip for 1 h. Unbound protein was rinsed off thoroughly, and the sample was imaged in a 5 mM Tris buffer. SpeedyTrack was performed with a vertical shift time of 7.5 μs for 15 rows after each exposure. (**a-c**) Results under a fixed excitation power of 280 mW (~22 kW/cm$^2$) and varied exposure times. (**a**) Example distributions of photon count per exposure at 50, 100, and 200 μs exposure times. (**b**) Mean photon counts per localization as a function of the exposure time. (**c**) Localization errors at different exposure times, determined by repeated detection of the same molecule in the SpeedyTrack data. (**d-f**) Results under a fixed exposure time of 300 μs and varied excitation laser powers. (**d**) Photon count per localization and background photons per pixel as a function of the excitation power (powers of 40-360 mW correspond to power densities of ~3-28 kW/cm$^2$). (**e**) Measured localization error $\sigma$ as a function of the excitation power. (**f**) Measured localization error in (e) versus that expected from Mortensen et al. (ref. 31), $\sigma_0$, based on the detected single-molecule and background photon counts in (d) for the different excitation powers. Red line: $\sigma_0$ scaled by $\sqrt{2}$ to account for the multiplicative noise intrinsic to EM-CCDs. Each data point in (b-f) is calculated from 3,000-15,000 single-molecule trajectories. Errors are smaller than the symbol size.

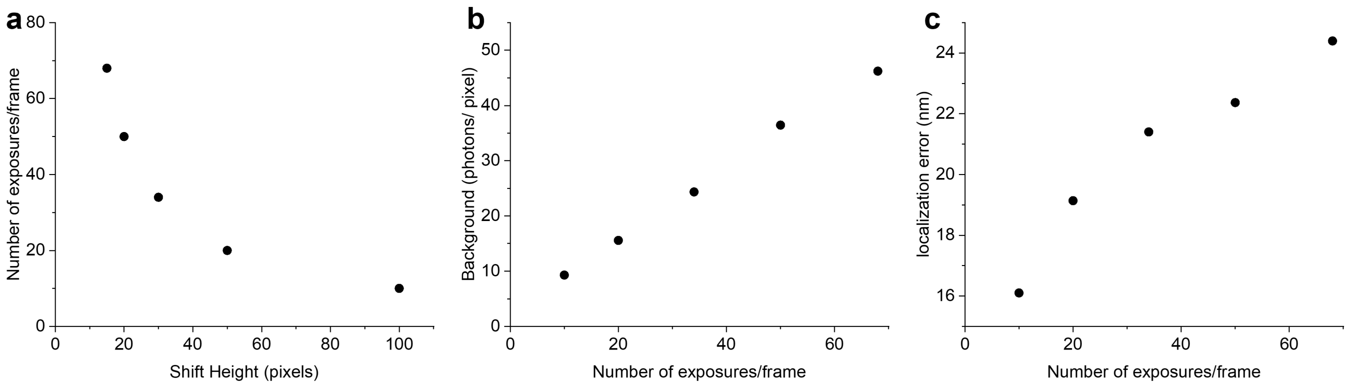


**Supplementary Fig. 4.** Impact of the shift height and accumulated background on SpeedyTrack results. (**a**) Number of exposures per frame as a function of shift height, namely, the number of rows shifted after each exposure. (**b**) Background in the SpeedyTrack image as a function of the number of exposures per frame, for detecting immobilized Cy3B-labelled avidin under the excitation power of 280 mW (~22 kW/cm$^2$) and 0.3 ms exposure time. (**c**) Localization error measured with different numbers of exposures per frame.

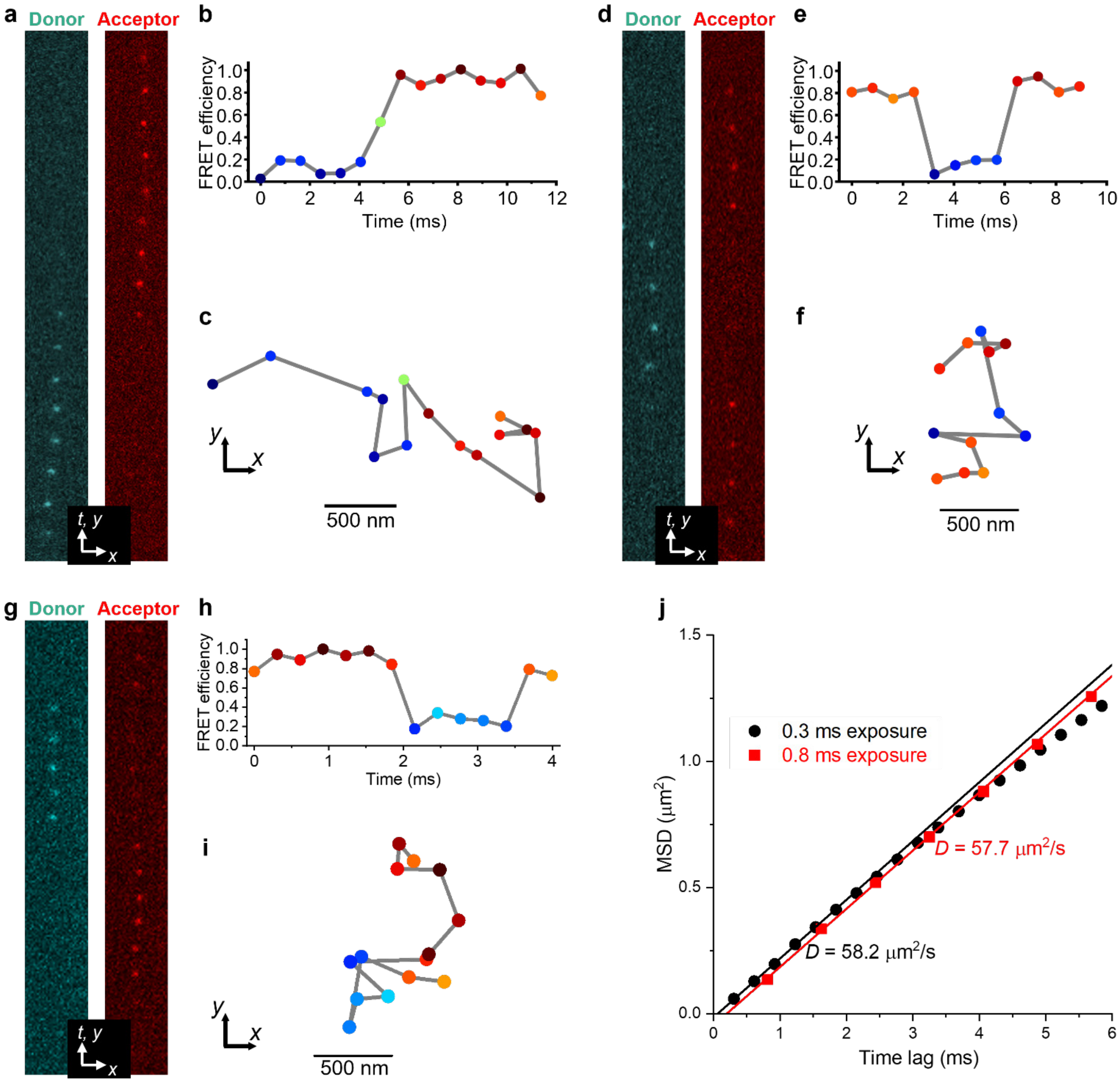


**Supplementary Fig. 5.** Additional SpeedyTrack smFRET data of freely diffusing dual-labeled DNA hairpins. (**a-f**) Additional example results acquired at an exposure time of 800 μs and a vertical shift time of 12.5 μs for 25 rows. (**a**) An example SpeedyTrack streak in the donor and acceptor channels for a hairpin molecule that switches from a low-FRET to a high-FRET state. (**b,c**) FRET efficiency time trace and reconstructed spatial trajectory from (a), colored by the FRET value. (**d-f**) Similar to (a-c), but for another molecule that switched back and forth between high-FRET and low-FRET states. (**g-i**) Example data acquired at an exposure time of 300 μs and a vertical shift time of 7.5 μs for 15 rows. This molecule also switched back and forth between high-FRET and low-FRET states (**j**) MSD vs. time lag calculated from trajectories obtained with 300 μs and 800 μs exposure times. Lines: Linear fits to the first 4 data points, yielding diffusion coefficients of 58.2 and 57.7 $\mu m^2/s$, respectively.

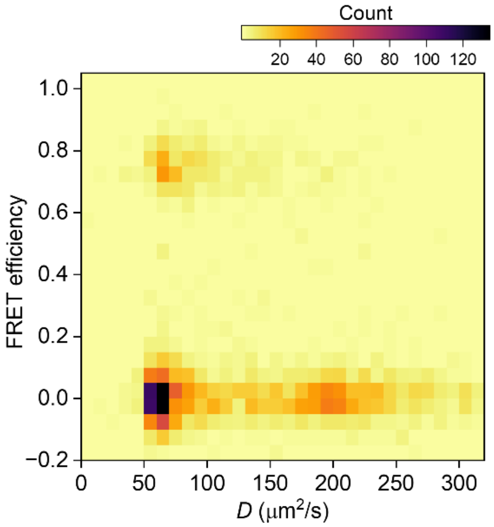


**Supplementary Fig. 6.** Two-dimensional distribution of mean FRET efficiency vs. estimated diffusion coefficient like that shown in Fig. 3h, but for SpeedyTrack results obtained with an exposure time of 50 μs.

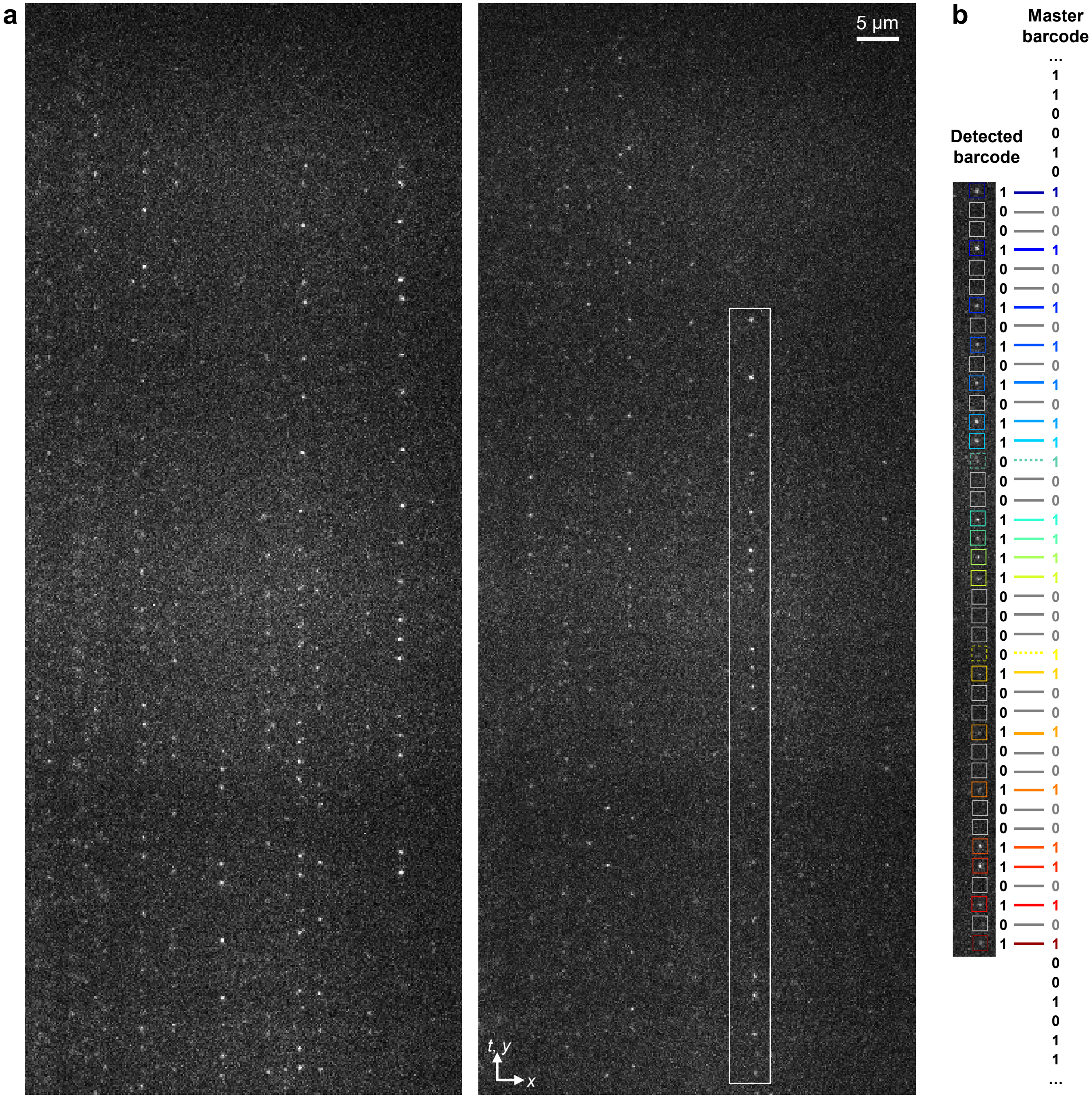


**Supplementary Fig. 7**. (**a**) Two example VS-SpeedyTrack frames of Dendra2 FP diffusing in the ER lumen of a living COS-7 cell, at 500 μs time resolution. (**b**) Aligning the boxed single-molecule streak in (a) with the master sequence. The best match is achieved with two missed localizations marked by the dashed boxes (thus a Hamming distance of 2), underlined in the master barcode as (reading from the bottom to top as 1s for detected single-molecule images and 0s for gaps): 10001001101001010110010010011000111100111010101001001010011001011011. The absolute vertical positions of the molecule at different timepoints are thus recovered by deducting the matched patterned vertical shifts in the sequence.

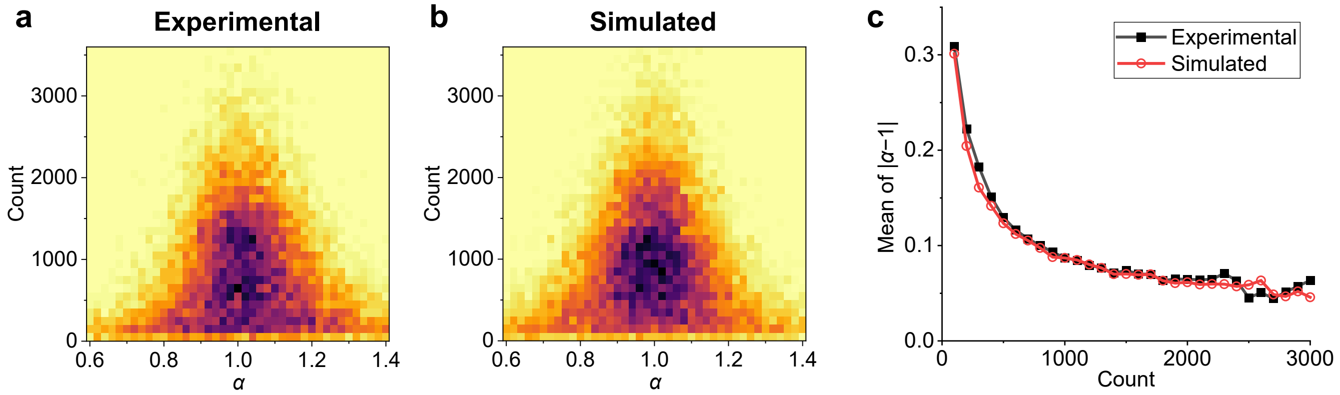


**Supplementary Fig. 8**. Comparison of the anomalous exponent *α* vs. count of displacements from experimental and simulated trajectories. (**a**) Two-dimensional distribution of *α* vs. count of displacements in individual spatial bins, as shown in the inset of Fig. 4j for the experimental VS-SpeedyTrack data of Dendra2 FP in the COS-7 cell ER lumen. (**b**) Distribution based on identical analysis of simulated trajectories. Trajectories were simulated under one-dimensional normal diffusion at $D$ = 9 μm$^2$/s with a timestep of 500 μs and a trajectory length distribution matching the experimental data. (**c**) Average absolute deviation of *α* from 1 as a function of count of displacements, for the experimental (black) and simulated (red) data.

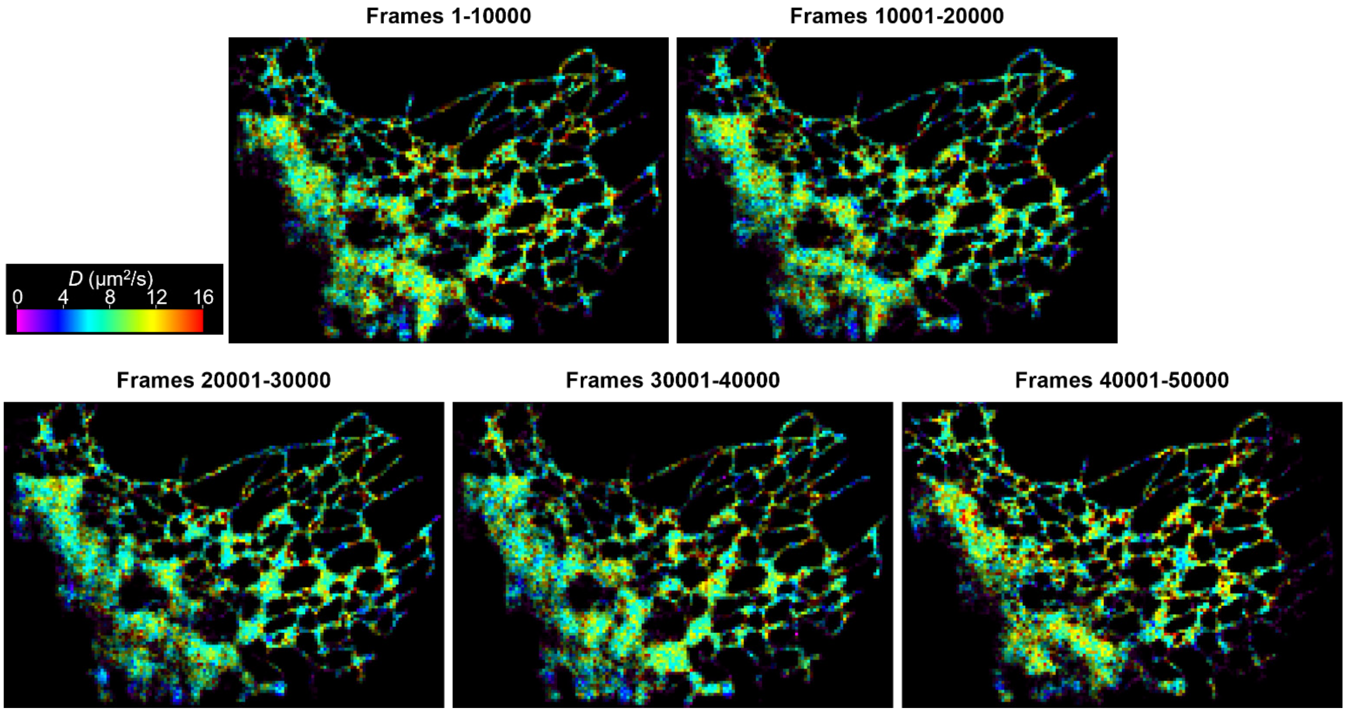


**Supplementary Fig. 9**. Color-coded $D$ maps of Dendra2 in the ER, constructed like Fig. 4h but after segmenting the dataset by frames, so that each $D$ map is constructed from 10,000 frames of VS-SpeedyTrack data acquired in ~15 min. A larger spatial grid size of 240×240 nm$^2$ is used to fit and color-render local diffusivity.

### Description of Supplementary Videos

**Supplementary Video 1**. Example SpeedyTrack data frames of Cy3B-labeled carbonic anhydrase diffusing in PBS, with an exposure time of 300 µs and vertical shift time of 7.5 µs for 15 rows at each timepoint. Left, center, and right panels show SpeedyTrack frames recorded at the beginning, middle, and end of one run, with frame numbers marked at the top.

**Supplementary Video 2**. Example VS-SpeedyTrack data frames of Dendra2 FP diffusing in the ER lumen of a living COS-7 cell, performed at 500 µs resolution. Left, center, and right panels show VS-SpeedyTrack frames recorded at the beginning, middle, and end of one run, with frame numbers marked at the top.
